# Mutation in *Wdr45* leads to early motor dysfunction and widespread aberrant axon terminals in a beta-propeller protein associated neurodegeneration (BPAN) patient-inspired mouse model

**DOI:** 10.1101/2024.12.13.628412

**Authors:** Brandon L Meyerink, Krishna S Karia, Mitchell J Rechtzigel, Prithvi R Patthi, Ariana C Edwards, Jessica M Howard, Elizabeth R Aaseng, Jill M Weimer, Louis-Jan Pilaz

## Abstract

Beta-propeller Protein Associated Neurodegeneration (BPAN) is a devastating neurodevelopmental and neurodegenerative disease linked to variants in *WDR45*. Currently, there is no cure or disease altering treatment for this disease. This is, in part, due to a lack of insight into early phenotypes of BPAN progression and *WDR45*’s role in establishing and maintaining neurological function. Here we generated and characterized a mouse model bearing a c52C>T BPAN patient variant in *Wdr45.* We show this mutation ablates WDR45 protein expression and alters autophagy in the brain. Behavioral analysis of these mice revealed characteristic signs of BPAN including cognitive impairment, hyperactivity, and motor decline. We show these behaviors coincide with widespread neuroinflammation and development of axonal spheroids in multiple neuron subclasses throughout the brain. Several lines of evidence suggest these spheroids arise from axon terminals. Transcriptomic analysis uncovered multiple disrupted pathways in the cortex including genes associated with synapses, neurites, endosomes, endoplasmic reticulum, and ferroptosis. This is supported by accumulation of the iron regulating transferrin receptor 1 (TFRC) and the endoplasmic reticulum resident calreticulin (CALR) in the cortex as these animals age. CALR forms spheroid structures similar to the axonal spheroids seen in these animals. Taken together, our data demonstrate that WDR45 is necessary for healthy brain function and maintenance of axon terminals. This model opens the door to therapeutics targeting BPAN and further exploration of the role of WDR45 in neuronal function.

## 1 Introduction

BPAN is a devastating neurodegenerative disease linked to mutations in *WDR45* located on the X chromosome (Haack et al., 2013, Hayflick et al., 2013). The disease onsets in infancy with seizures and developmental delay followed by attention deficit, sleep disruptions, and intellectual deficiency in childhood. Clinical imaging of BPAN patients shows decreased volume of the white matter tracts which resolves by unknown mechanisms before adolescence (Adang et al., 2020, Papandreou et al., 2022). In early adulthood, seizures decline though patients develop parkinsonism, dystonia, and progressive dementia (Hayflick et al., 2013, Wilson et al., 2021). At this late stage, iron accumulates in the substantia nigra and the basal ganglia though this accumulation does not necessarily correlate with the onset or severity of other symptoms (Wilson et al., 2021). Post-mortem histology reveals significant neurodegeneration in the cortex, cerebellum, locus coeruleus, and substantia nigra (Paudel et al., 2015). Ultimately, BPAN is lethal, often in mid-adulthood (Hayflick et al., 2013, Saitsu et al., 2013, Adang et al., 2020). Most BPAN patients have a single mutation with heterozygous expression of the *WDR45* gene. It is thought that in this population, X-inactivation leads to expression of wildtype and affected alleles in distinct cell populations throughout the body, though the clinical signatures of BPAN are entirely neurological. Males represent a small portion of the patient population and often present more severe disease progression (Adang et al., 2020). There is currently no disease altering treatment for BPAN, only symptomatic relief with drugs targeted at movement and seizure symptoms (Wilson et al., 2021). Crucially, we lack an understanding of WDR45 function, especially in the central nervous system (CNS), which is needed for the development of targeted therapies. WDR45’s sequence is well conserved, with mice and humans sharing nearly 98% homology (Uniprot), thus making mice an enticing model for understanding protein function and disease pathobiology. Several WDR45 knockout mouse models have been established to answer these questions. The first model ablating WDR45 used a floxed allele in conjunction with a Nestin-Cre driver that conditionally removes *Wdr45* expression in the CNS (Zhao et al., 2015). These animals exhibit clear neurological deficits including memory and learning deficiencies, motor decline, and neuronal loss, albeit at an advanced age of 13-months. Interestingly, eosinophilic spheroids appear in Purkinje neuron projections to the deep cerebellar nuclei at 1.5-months. These animals also accumulate telltale signs of autophagy dysfunction with increased LC3 and p62 protein levels. Subsequently, two full body *Wdr45* knockout animals were generated using CRISPR or TALEN technologies (Wan et al., 2020, Biagosch et al., 2021). These animals presented with similar phenotypes compared to the first model. In addition, however, these newer models displayed earlier signs of neurological dysfunction including increased seizure susceptibility, social avoidance, anxiety phenotypes as well as retinal thinning, and auditory decline in later stages. Of note, many of these phenotypes were only reported in complete knockout animals (hemizygous mutation for males, homozygous mutation for females) and not in heterozygous females, which represent the main patient populations (Zhao et al., 2015, Biagosch et al., 2021). Collectively, these studies demonstrate that mice lacking *Wdr45* expression display many phenotypes analogous to BPAN though they appear much later than the equivalent stages in patient populations.

Important questions remain about the impact of WDR45 loss in the brain. Primarily, the link between early cellular and molecular insults and neuronal dysfunction at various stages of the disease is not clear, owing primarily to the paucity of early pathological changes in the current rodent models. Knowing these early phenotypes will allow us to define critical periods delineating cause from effect and defining windows for therapeutic intervention.

Here we characterized a novel mouse model of BPAN bearing the c52C>T patient variant, which is predicted to ablate all but the first few amino acids of the protein and is the first model to directly mimic a patient mutation (ClinVar). We performed longitudinal assessment of behavioral phenotypes and pathological changes to generate a roadmap of disease development. This innovative model exhibits earlier signs of neurological deficits and a wider variety of axonal degeneration than previously reported in other models. We further demonstrate that heterozygous mutant mice, directly reflecting the majority of the patient population, exhibit robust behavioral and pathological phenotypes. These findings are critical to understanding how BPAN develops and establishing key markers of disease progression, paving the way for future therapeutic testing and studies of the molecular role of WDR45 in building and maintaining proper neurological function.

## 2 Materials and methods

### 2.1 Animals

All experiments were performed in accordance with Sanford Research IACUC guidelines. All animals in these experiments were on the C57BL/6J background with mice purchased from Jackson Laboratory and maintained as a breeder colony. During behavioral experiments, experimenters were blinded to animal genotypes.

### 2.2 CRISPR based mouse generation

Guide RNA and homology directed repair templates for CRISPR editing were designed using the Alt-R CRISPR Design Tool hosted by IDT. Oligonucleotides and CAS9 were ordered from Integrated DNA Technologies (IDT) using the Alt-R design tools and reagents (sequences and ordering information in Table S1). The transactivating CRISPR RNA (tracrRNA) and targeted CRISPR RNA (crRNA) were hybridized in a 1:1 molar solution at 98 °C for 2 minutes then held at room temperature for 10 minutes, resulting in the final guide RNA (gRNA). The CRISPR working solution (30 µM gRNA, 50 µM single stranded repair template, 1mg/ul Cas9 protein, .02% fast green dye in Opti-MEM) was incubated at 37° C for 10 minutes prior to surgery.

Small animal surgery to introduce CRISPR reagents into the nascent embryo followed the improved Genome editing via Oviductal Nucleic Acids Delivery (iGONAD) protocol (Ohtsuka et al., 2018, Gurumurthy et al., 2019). Briefly, embryonic day 0.7 pregnant dams were anesthetized and maintained using isoflurane. Under aseptic conditions, hair was removed from the dorsal abdomen and an incision was made on the dorsal lateral portion of the abdomen. The distal uterine horn including ovary and oviduct were exposed from the dorsal abdominal cavity. ∼1µL of the CRISPR working solution was injected into the distal oviduct using a pulled glass capillary needle. The oviduct was then covered with a small square of sterile Kimwipe and wetted using sterile saline before electroporation using a tweezer electrode and Bex pulse generator (CUY21EDIT II). Electroporation consisted of 3 square wave pulses set at 5 ms, 60V, 125 mA with 10% decay. Following electroporation, the Kimwipe was removed, the uterus was replaced into the abdomen, and the procedure was repeated for the contralateral side. The muscle and skin layers were then sutured closed. Animals were allowed to recover from anesthesia under observation and given ad libitum Ketorolac for pain management.

### 2.3 Genotyping of animals

Primers were designed to flank the region covered by the homology directed repair template. These primers were used to generate an amplicon that was then sent for sanger sequencing of the amplified region (Eurofins) to confirm the desired mutation and ensure there were no undesired indels in the target region. This was performed for F0 and F1 animals from the iGONAD surgeries that were used to generate study animals. For subsequent generations, a Taqman genotyping probe was purchased from Thermo Scientific and used for genotyping (Assay ID: ANWDDVT).

### 2.4 Force-plate actimeter

Individual mice were placed in a force-plate actimeter (BASi, West Lafayette, IN) for 20 minutes each. The forceplate was positioned in a sound attenuating box to reduce outside interference with animal behavior. Tracking and analysis was performed using the FPA analysis software.

### 2.5 Perinatal vocalization

Perinatal vocalizations were recorded at perinatal day (P) 5, 7, and 10 based on previously described protocols (Moy et al., 2009, Shekel et al., 2021). Each testing day, pups were removed from the dam and placed in the home cage on a heating pad for 5 minutes prior to introduction to the recording apparatus. The recording apparatus consists of a sound attenuating box with a glass beaker to ensure the animal is well positioned for recording. An ultrasonic vocalization (USV) detecting microphone (Avisoft Bioacoustics) was placed through the top of the box facing the opening of the glass beaker. We then placed the animal into the glass beaker for 5 minutes and recorded any ultrasonic vocalizations using the Avisoft-Recorder software. We then returned the animal to the dam after the 5-minute recording period. Spectrograms were analyzed using the Avisoft SASlab-Pro software.

### 2.6 Morris Water Maze

We performed the Morris water maze analysis over a 5 day period in a circular pool (1.2 meter diameter) filled to a depth of 66cm with 23°C water (Johnson et al., 2023). Four unique cues were placed on the interior pool wall above the water level at 0, 90, 180, and 270 degrees. On the morning of the first day, probe test training was performed where the goal platform is placed under clear water and a flag is placed on the platform. Mice were then placed in one of the four quadrants of the apparatus and allow them to explore the apparatus for 60 seconds. After this time, if the mice had not found the platform, they were guided to the platform by the experimenter. Mice were allowed to sit on the platform for 20 seconds before the next trial to reinforce learning. This same procedure was followed by starting the mouse in the three other quadrants. The mice were then given a three-hour rest period before they were subjected to the next round of trials in the afternoon following the same procedure. For the next four days, we subjected the animals to test trials. These test trials were performed in the same apparatus though without the flag marking the platform and with gothic white, non-toxic, liquid tempura paint used to make the water opaque. Mice were started in all four positions of the apparatus for both morning and afternoon sessions of this testing. Time to reach platform and swim speed during the trials were assessed using the ANY-maze software, which tracks animal movements.

### 2.7 Marble Bury

Marble bury behaviors were assessed using an established protocol (Angoa-Perez et al., 2013). A single mouse was placed in a clean cage where 12 marbles of equal size sit on top of the bedding in a standard pattern (three columns of four marbles). The cage was then enclosed, and the mice were allowed to free roam the cage for 30 minutes. After this time the mice were removed from the cage and replaced into their home cage. Marbles were assessed and any marbles buried at least 2/3^rd^ of the way are counted as buried.

### 2.8 Accelerating Rotarod

Mice were placed on the rotarod device (AccuScan Instruments, Inc.) at 0 rpm (Kovacs and Pearce, 2015, Poppens et al., 2019). The rotarod then began slowly accelerating by 0.2 rpm / second. This acceleration continues for 4 minutes or until all mice have fallen off the apparatus. The time between the start of acceleration and when the animal falls below the rotarod is noted. Each trial consists of 3 consecutive runs on the rotarod for each mouse. The mice are first subjected to a training trial followed by a 90-minute rest period. The mice were then subjected to three trials with a 30-minute rest period between each trial. The average time to fall for these trials was used for our analysis.

### 2.9 Vertical Pole Climb

Vertical pole climb testing was performed using a vertical, threaded rod (diameter = 1.27cm, height = 60 cm) attached to a large metal base (Johnson et al., 2021). The base was covered with padding to ensure the safety of the animal. The animal was placed, head facing downward, at the top of the vertical rod. Once the animal is placed, the time taken for the animal to climb down the threaded rod and reach the padded base is noted. If the animal had not reached the base after 60 seconds, the animal was removed from the rod and given a time of 60 seconds for that trial. This procedure was repeated four more times for each animal with a 60 second rest period between each trial.

### 2.10 Antibodies

Antibodies for these experiments are outlined in Table S1.

### 2.11 Tissue collection and preparation

Animals were euthanized via CO2 inhalation followed by cardiac perfusion using ∼6ml of Phosphate Buffered Saline (PBS). The brain was then removed and hemisected with the right hemisphere placed in a tube on dry ice for molecular assays. The left hemisphere was placed in 4% paraformaldehyde in PBS and stored overnight at 4°C. After fixation, the tissue was moved to PBS with 0.02% Sodium Azide and stored 4°C.

### 2.12 Immunohistochemistry

Fixed brains were sagittally sectioned via vibratome at 50μm and stored in Tris Buffered Saline (TBS) (Cain et al., 2019). Tissue sections were then transferred to 0.1% hydrogen peroxide for 10 minutes then washed 3 in TBS times for 5 minutes. These sections were transferred to blocking solution (TBS, 10% Goat Serum, 0.3% Triton X-100) for 30 minutes with agitation. They were then placed in fresh blocking solution with diluted primary antibodies overnight at 4°C with agitation. The next day, tissues were washed 3 times with TBS for 5 minutes each in TBS before adding blocking solution with biotinylated secondary antibodies and incubating at room temperature for 2 hours. Tissues were then washed 3 times with TBS before the addition of VECTASTAIN ABC-HRP reagent diluted in TBS based on manufacturers’ recommendations. Tissues were incubated for 1 hour followed by another 3 washes in TBS. Finally, 3,3′-diaminobenzidine was dissolved in TBS and filtered through a 0.45μm syringe filter before tissues incubation. Tissues were allowed to incubate until a dark stain appeared under the microscope and the reaction was quickly stopped with the addition of cold TBS. The tissues were then washed 3 times with cold TBS. Tissues were then mounted on charged slides and dehydrated through air-drying followed by an alcohol gradient. Finally, tissues were covered using DPX mounting medium and coverslipped. These slides were allowed to be cured overnight before being imaged on using the Aperio AT2 slide scanner (Leica) using a 20x objective.

### 2.13 Immunofluorescence

Immunofluorescence was performed on 50μm sagittal sections. Tissue sections were then transferred to blocking solution (TBS, 10% Goat Serum, 0.3% Triton X-100) for 30 minutes with agitation. They were then placed in fresh blocking solution with diluted primary antibodies overnight at 4°C with agitation. The next day, tissues were washed 3 times with TBS for 5 minutes each before adding blocking solution with secondary antibodies. Tissues were incubated with secondary antibodies at room temperature for 2 hours before being washed 3 times with TBS. Tissues were then mounted on charged slides and cover slipped using DAKO faramount aqueous mounting medium and allowed to cure overnight before being imaged using either the Nikon 90i epifluorescent microscope or the Nikon A1 SORA confocal microscope with 20x or 60x objectives.

### 2.14 Western Blotting

Western blotting was performed on cortical brain homogenates lysed using RIPA buffer. Concentration of the lysates was determined using the Thermo Fisher BCA colorimetric kit and 20μg of each protein sample was mixed with loading buffer (Thermo Fisher) diluted to working concentration and denatured 70°C for 10 minutes prior to loading on 4–20% Mini-PROTEAN TGX Stain-Free Protein Gels (Bio-rad) or 4-12% Bolt Bis-Tris Plus Mini Protein Gels (Invitrogen). Proteins were transferred to nitrocellulose membranes, imaged for stain free protein quantification, and incubated in 5% milk dissolved in TBS with 0.1% Tween 20 (TBS-T).

Membranes were then incubated in primary antibodies overnight at 4°C, washed with TBS-T and incubated with HRP bound secondary antibodies diluted in TBS-T for 2 hours prior to final washes in TBS-T, TBS and incubations in Pierce ECL Western Blotting Substrate (Thermo Fisher) before imaging on a Chemi-MP imager (Bio-Rad). Bands were quantified by densitometry using ImageLab (Bio-Rad) and normalized to the protein loading assessed by the stain-free total protein of each lane. Membranes were then stripped of antibodies using Restore Plus stripping buffer (Thermo) for 20 minutes then washed 3 times in TBS. These membranes were reimaged to ensure the loss of signal before being reblocked and reprobed using the above protocol.

### 2.15 RNA extraction, RT-qPCR. and Sequencing

RNA was extracted from mouse cortices using a Trizol reaction. Tissue was homogenized using Trizol reagent prior to adding chloroform to the homogenized sample. Samples were then centrifuged in this solution to fractionate and the top clear fraction was removed. This fraction was then added to isopropanol and centrifuged again to pellet out the RNA from the sample. Ethanol was then added to resuspend the RNA pellet before further centrifugation. We then removed the ethanol from the tube and allowed the pellet to dry prior to resuspending the RNA pellet in molecular biology grade water. RNA was then converted to cDNA using the iScript cDNA synthesis kit (Bio-Rad) following manufacturer’s instructions.

RNA sequencing was performed by Novogene. Sequencing files were downloaded from the Novogene server and uploaded to Galaxy (usegalaxy.org) for analysis. Forward and reverse read files were paired and adaptors were trimmed using Cutadapt. Reads were then mapped to the reference genome (mm10) using RNAstar. Read counts were determined using featureCounts and the differentially expressed genes were assessed using DESeq2. Results were analyzed and differentially expressed genes, defined by p < 0.05, were used for ontology and pathway analysis. Genes with positive and negative fold changes were separately input into Enrichr (https://maayanlab.cloud/Enrichr/) and significant ontologies and pathways were downloaded. This tool was accessed on 5/15/24.

cDNA from 1-month wildtype and Wdr45c52C>T males was isolated from cortical samples as described above. Primers were designed against Wdr45 and Gapdh (sequences in Table S1) and RT-qPCR was performed using qPCR Master Mix with SYBR Green (Gold Bio) per manufacturer’s instructions. Signal was normalized to GAPDH signal for each sample.

### 2.16 Statistical methods

Statistical analysis was performed using GraphPad Prism 9. Details of the individual tests used are noted in the figure legends. T-testing was used for comparing two groups. One-way ANOVA with Holm-Šídák post hoc test was used for three compared groups. Two-way ANOVA with Holm-Šídák post hoc test was used for longitudinal behavior analysis. The Mantel-Cox test was used for survival curve testing. Values represent the mean ± SEM. For individual timepoints, * = p<0.05, ** = p<0.01, *** = p<0.001, **** = p<0.0001. # are used to show genotype effects from ANOVA. # = p<0.05, ## = p<0.01, ### = p<0.001, #### = p<0.0001.

## 3. Results

### 3.1 Wdr45 c52C>T hemizygous and homozygous mice lack WDR45 protein expression

We introduced a mutation in the *Wdr45* genes via CRISPR editing using the iGONAD electroporation technique in C57BL/6 J mice (Ohtsuka et al., 2018, Gurumurthy et al., 2019). The induced mutation targets the first coding exon of *Wdr45* mimicking a patient variant (Figure 1A) (ClinVar). This single nucleotide mutation lays in a conserved region of the gene and introduces a premature stop codon early in the coding region. If translated, the resulting peptide would retain only 18 amino acids. After CRISPR editing, mice were backcrossed to wild type C57BL/6 J mice for at least two generations. Sequencing of the target region indicated no unwanted indels or mutations (Figure 1B-C). The later generations were bred to produce the following genotypes: male *Wdr45 -/Y* (hemizygous), female *Wdr45 +/-* (heterozygous), and female *Wdr45 -/-* (homozygous) mice as well as the control *Wdr45* +/+ (wildtype) mice that were used for these studies. We observed no decrease in *Wdr45* transcript measured through RT-qPCR and next-gen RNA sequencing in cortical samples (Figure S1B). This suggests *Wdr45* transcript may not undergo canonical nonsense-mediated decay with this mutation. However, as predicted, this mutation completely ablated full length WDR45 protein expression in hemizygous and homozygous (knockout) cortices and significantly reduced expression in heterozygous cortices as indicated by western blotting (Figure 2B,D,K,M). *Wdr45* c52C>T mice did not die earlier than their wildtype littermates (Figure S1C) and only hemizygous males showed a decrease in body weight compared to wildtype littermates (Figure 3F-G). Together, these data demonstrate successful generation of a single nucleotide mutation in *Wdr45* mimicking a patient variant which leads to loss of full length WDR45 expression.

**Figure 1:**
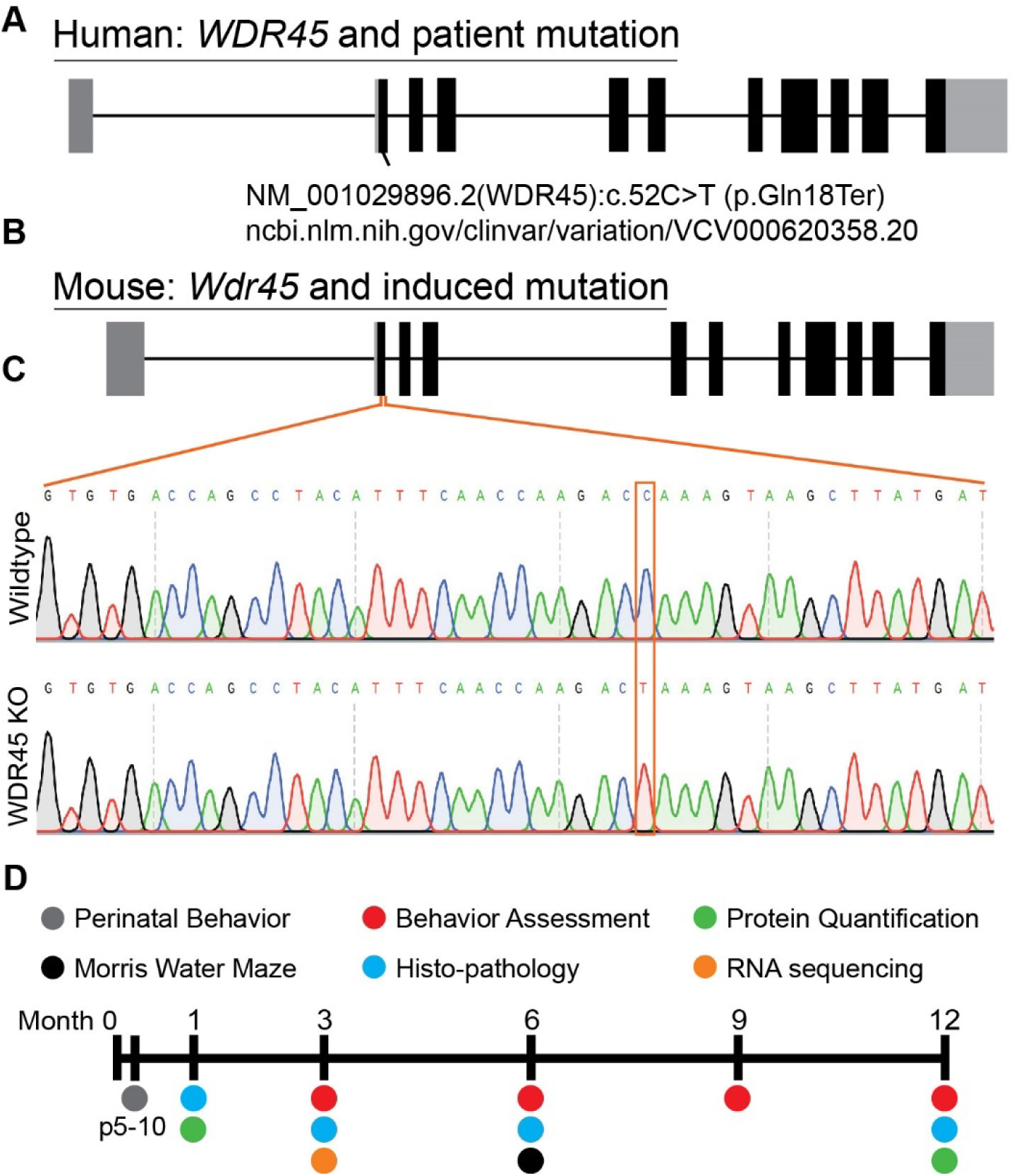
Mutation of mouse *Wdr45* to mimic BPAN patient variant. Mutations in human *WDR45* are associated with the disease BPAN. Our mouse model was inspired by the c52C>T patient mutation (A,B). Sanger sequencing of the region surrounding the mutation confirmed introduction of the target mutation (C). Timeline of experiments in this study (D).

**Figure 2:**
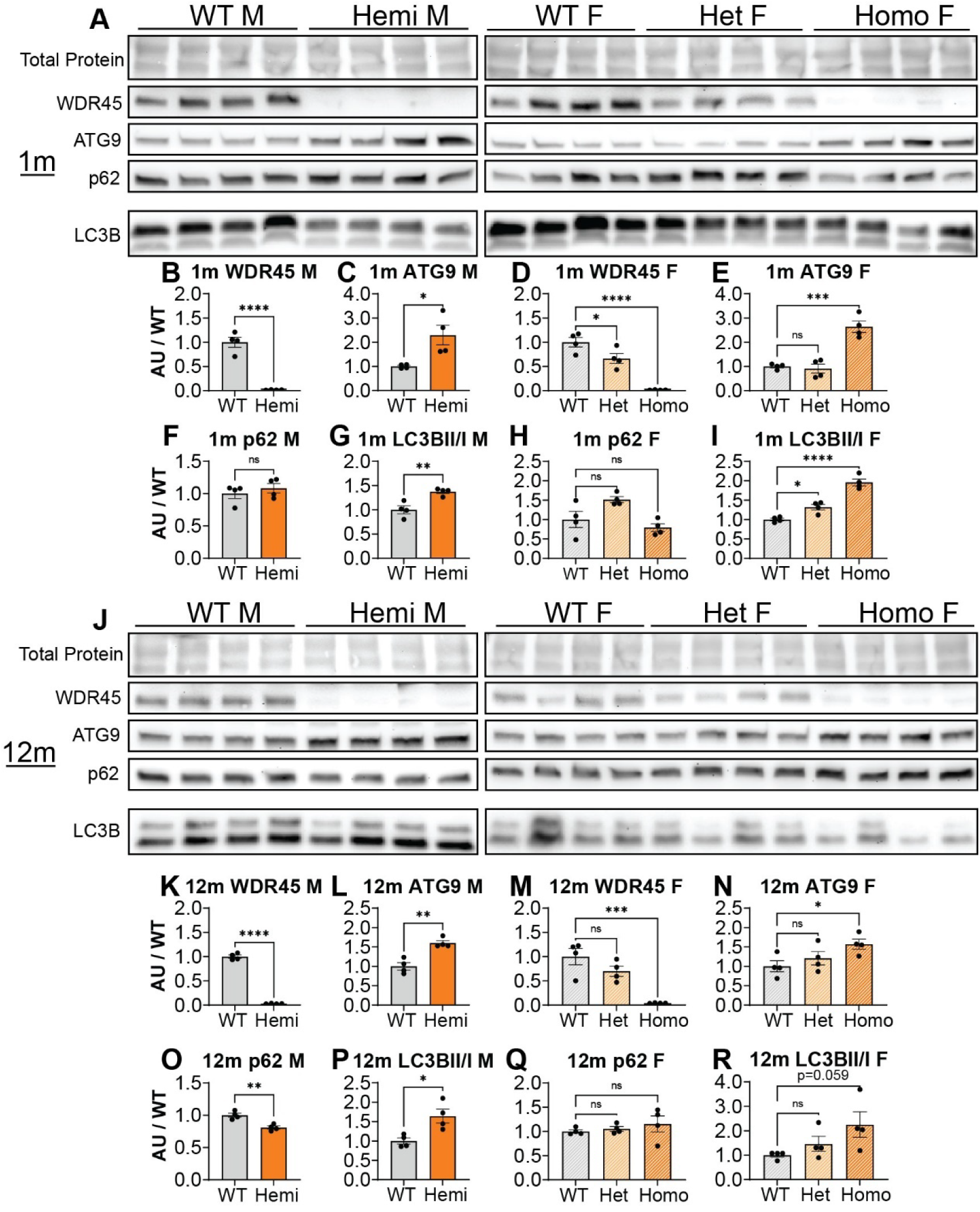
*Wdr45* c52C>T mutation ablates full-length WDR45 expression and alters autophagy protein levels in cortical samples. Western blotting was performed on cortical samples from 1-month (A) and 12-month (J) animals. Western blotting assay shows complete loss of full-length WDR45 protein (B,D,K,M). ATG9 (C,E,L,N) levels were altered at both timepoints and LC3B II / I ratio (G,I,P,R) was altered in *Wdr45* c52C>T animals at 1-month and in males at 12-months. WDR45, ATG9, and p62 signal was normalized to total protein for each lane. For Males: T-test was used. For Females: One-way ANOVA was used with Holm-Šídák post hoc test, N=4 animals for each group. Mean ± SEM. * = p<0.05, ** = p<0.01, *** = p<0.001, **** = p<0.0001.

**Figure 3:**
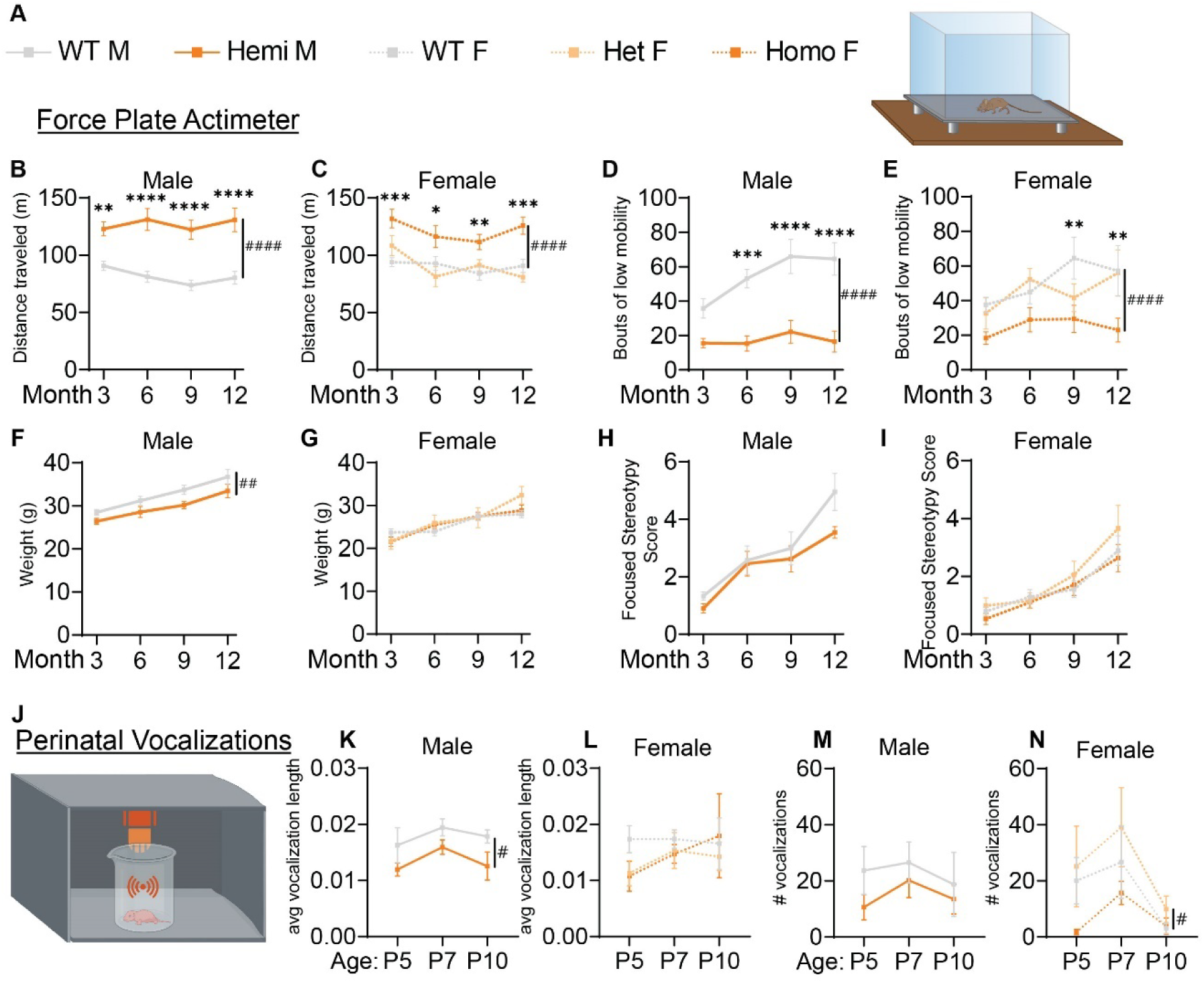
*Wdr45* c52C>T mutation in mice leads to hyperactivity and altered perinatal vocalizations. Genotype key for line graphs (A). Forceplate actimeter assay showed more movement of the *Wdr45* c52C>T mice during the testing period (B,C) and fewer bouts of low mobility (D,E), but no stereotypy associated with increased anxiety (H,I). Hemizygous males showed a decreased body weight over the entire testing period (F). Heterozygous and homozygous females showed no change in body weight compared to wiltype (G). Ultrasonic vocalization analysis was performed on perinatal animals (J). Average length of individual vocalizations (K,L) and the number of vocalizations during the recording period (M,N) were analyzed. Two-way ANOVA was used with Holm-Šídák post hoc test. Mean ± SEM. * = p<0.05, ** = p<0.01, *** = p<0.001, **** = p<0.0001. # are used to show genotype effects from ANOVA. # = p<0.05, ## = p<0.01, ### = p<0.001, #### = p<0.0001.

### 3.2 Wdr45 c52C>T mice show signs of altered autophagy

WDR45 has previously been implicated in autophagy and particularly in the development of the autophagosome (Bakula et al., 2017). Thus we sought to quantify the abundance of markers for this pathway in their novel *c52 C>T* model at 1- and 12-months of age. Upon its integration into the autophagosome membrane, free LC3B (LC3B-I) is cleaved and lipidated, thus leading to the generation of another protein species (LC3B-II) observed by western blot (Tanida et al., 2008). ATG9 is thought to decorate the membrane of small vesicles used as seeds for autophagosome biogenesis (Holzer et al., 2024). P62 labels proteins bound for degradation via autophagy. Thus, any difference in the ratio between the two LC3B species, as well as ATG9 and P62 protein levels can be used to highlight defects in the autophagy pathway (Mizushima et al., 2010). While P62 did not consistently show altered levels, LC3B-II / LC3B-I ratios and ATG9 levels were higher in the cortices of hemizygous and homozygous *Wdr45 c52C>T* animals compared to their sex-matched littermates at both timepoints (Figure 2A, J). These data show that the autophagy pathway is indeed disrupted, though autophagic flux may not be altered in these animals.

### 3.3 *Wdr45 c52C>T* mice display increased locomotion

Given the early onset of BPAN in patients and progressive nature of the disease, we performed longitudinal behavioral assessments to examine progression of disease phenotypes. Force plate actimeter assessment was used to track movement, tremors, and movements termed stereotypy which often reflect excessive grooming and thus elevated anxiety and compulsive behaviors. Starting at 3-months, *Wdr45 c52C>T* animals show increased locomotion , traveling a longer distance during the allotted time and with fewer bouts of low mobility (Figure 3B-E). We did not see changes in the focused stereotypies of these animals suggesting this is not coincidental with an increase in anxiety-related grooming behavior (Figure 3H-I). However, this may be confounded by fewer bouts of low mobility leading to fewer opportunities to groom. This interpretation is supported by the frequent observation of their behavior in their home cage, as well as data from the marble bury test, commonly used to assess anxiety in mice (Jimenez Chavez and Szumlinski, 2024). This test showed no difference between genotypes (Figure S1D). These data indicate a hyperactive phenotype, a common finding in established models of neurological disorders and consistent with BPAN patient phenotypes (Walker et al., 2011, Jul et al., 2016, Wang et al., 2022, Wilson et al., 2021, Gregory et al., 2017).

### 3.4 Perinatal Wdr45 c52C>T mice have altered vocalization behavior

We next assessed the ultrasonic vocalization behavior of these animals in response to being placed away from their mother. These vocalizations are a critical way for neonatal mice to communicate with their mother and altered vocalization behavior is common in mouse models of developmental disorders including autism spectrum disorder (Moy et al., 2009, Shekel et al., 2021). We found that over the ages of P5, P7, and P10, hemizygous male mice had shorter ultrasonic calls whereas homozygous female mice had fewer calls overall than their wildtype littermates (Figure 3J-N). These data underscore that the neurological effect of loss of WDR45 is occurring very early in the development of these mice.

### 3.5 Wdr45 c52C>T mice exhibit learning and memory deficits

Along with the developmental delay, BPAN presents with intellectual disability especially later in life (Gregory et al., 2017, Hayflick et al., 2013). To test if this is recapitulated in *Wdr45 C>T* mice, we performed Morris water maze testing on mice at 6-months of age. During the probe training period, we found that hemizygous and homozygous animals were significantly less likely to find the platform (Figure 3D). Mice that were able to reach the platform in more than half the trials during the training period were then subjected to further testing over several days. We found hemizygous males took longer to reach the platform than wildtype littermates despite an increased swim speed (Figure 4C,E). However female mice showed no difference in time to platform, though this may be due to many of the homozygous females being excluded during the probe training period (8 of 13 homozygous animals). Those excluded were likely to have the most dramatic impairment in the task (Figure 4D). Together, these data indicate *Wdr45 c52C>T* mice have impaired performance in learning and memory testing consistent with cognitive impairment seen in BPAN patients.

**Figure 4:**
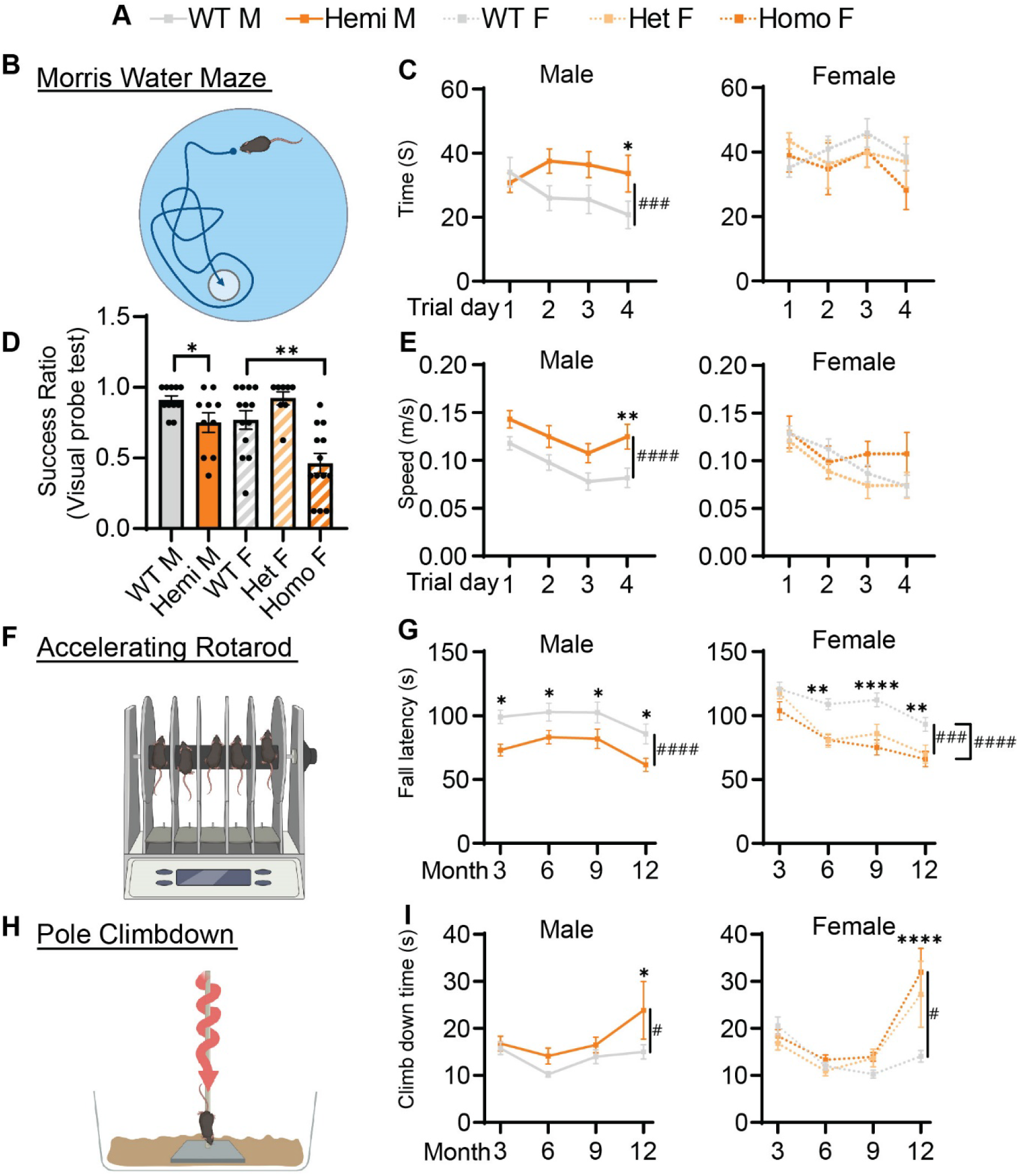
Cognitive and motor function of *Wdr45* c52C>T animals is impaired. Genotype key for line graphs (A). Morris water maze analysis (B) was performed on 6-month-old animals. Visual probe training showed reduced success in finding the platform for homozygous and heterozygous animals (D). Mice with success ratios above 0.5 were tested further with time to reach platform over the trials (C) and swim speed during trials (E) showing differences in hemizygous males. Rotarod analysis (F) showed fall latency was reduced for Hemizygous, Heterozygous, and Homozygous animals (G). Pole climbdown testing (H) showed mice at an advanced age took longer to climb down the pole (I). D used a T-test for males and a One-Way ANOVA with Holm-Šídák post hoc test for females, N=8-13 for each group. C,E,G,I used Two-way ANOVA with Holm-Šídák post hoc test, N=5-13 animals for each group. Mean ± SEM: * = p<0.05, ** = p<0.01, *** = p<0.001, **** = p<0.0001. # are used to show genotype effects from ANOVA. # = p<0.05, ## = p<0.01, ### = p<0.001, #### = p<0.0001.

### 3.6 Wdr45 c52C>T mice show motor defects

Motor decline is common in patients with BPAN especially in the later part of the disease (Hayflick et al., 2013). We tested if our BPAN mouse model has motor deficits using a rotarod assay accelerating linearly over time (Figure 4F). Hemizygous males fell off the rod earlier than their wildtype littermates as early as 3-months of age (Figure 4G). Homozygous and heterozygous females showed a similar pattern though this deficit beginning at 6-months of age (Figure 3G). Using a different motor assay, the pole climb-down assay, we tested the animal’s ability to climb down a vertical threaded pole in a given time (Figure 4H). *Wdr45 c52C>T* mice showed significant difficulty climbing down the rod at 12-months of age, indicating the motor defects are progressing around this timepoint (Figure 4I). These data show that *Wdr45 c52C>T* mice have motor deficits, starting relatively early in life and extending throughout the animal’s lifespan.

### 3.7 Dopaminergic and Noradrenergic neurons of Wdr45 c52 C>T mice display distal axon abnormalities

With the pathology seen in the substantia nigra of BPAN patients (Saitsu et al., 2013, Paudel et al., 2015), we evaluated the dopaminergic motor pathways using immunolabeling of Tyrosine Hydroxylase (TH), an enzyme critical for the production of dopamine, allowing us to visualize the dopaminergic neurons of the nigrostriatal tracts. At all ages examined, there were no observed changes in the density of the cell bodies or proximal axonal tracts of TH+ neurons residing in the substantia nigra (Figure 5B,C). However, starting at 3-months, there were large focal areas of TH+ immunoreactivity in the striatum which contains the distal ends of dopaminergic axons (Figure 5D). These appeared exclusively in the *Wdr45 c52C>T* animals, including heterozygous females (Figure 5 H-I). Indeed, we visualized large spheroids (approximately 5 microns in diameter) abundant in the caudate putamen and nucleus accumbens but completely absent closer to the substantia nigra and along the nigrostriatal fiber tracts (Figure 5C). These data indicate distal axons of dopaminergic neurons are distressed in *Wdr45 c52C>T* mice though the somatic and proximal projections appear unaffected.

**Figure 5:**
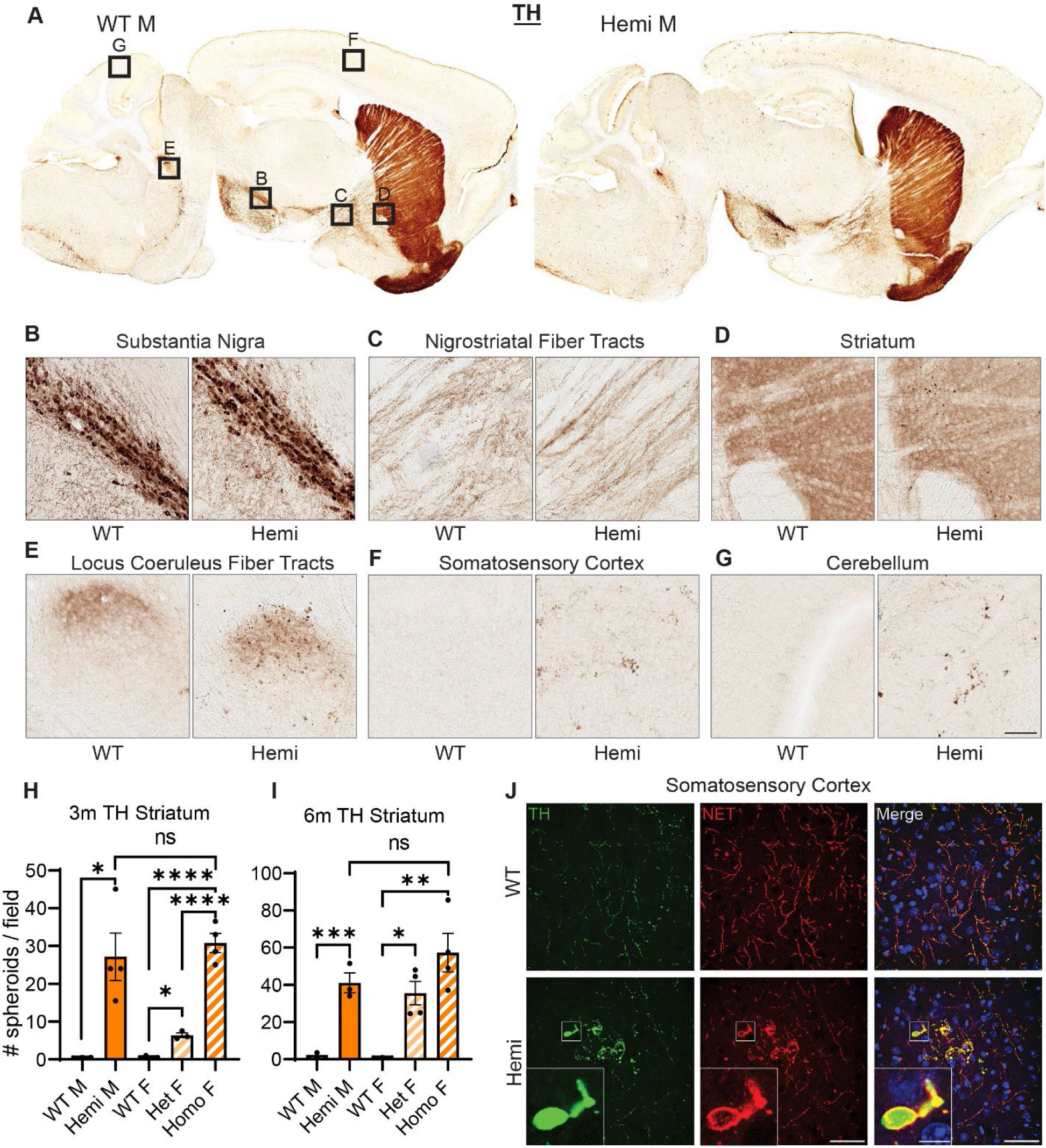
*Wdr45* c52C>T animals form large spheroids in the distal axons of Tyrosine Hydroxylase positive neurons. Immunohistochemistry labeling Tyrosine Hydroxylase (TH) was performed on brains from 12-month-old animals (A). Imaging in the Substantia Nigra (B) and the proximal nigrostriatal tracts (C) show little change in morphology in Wdr45 c52C>T animals. More distal TH positive innervation in Wdr45 c52C>T animals show large areas of immunostaining indicating a morphology change in these distal projections (D-G). Quantification of TH positive spheroids in the striatum at 3-months (H) and 6-months (I) of age. Immunofluorescent labeling of TH (Green) confirms these structures and colabeling of Norepinephrine Transporter (NET) (Red) indicates that these structures in the cortex are surrounded by this transmembrane transporter (J). Scale bars: B-G = 100µm, J = 50µm, J inset = 5µm. Individual field values were subjected to outlier analysis using the ROUT method (Q=1%). H,I used a T-test for males and a One-Way ANOVA for females with Holm-Šídák post hoc test, N=3-4 for each group. Mean ± SEM: * = p<0.05, ** = p<0.01, *** = p<0.001, **** = p<0.0001

Broadening our search for these axonal TH+ spheroids showed localization throughout the brain including the cortex, cerebellum, hippocampus, and within the projection tracts of the locus coeruleus (Figure 5E-G). The cerebellum and the locus coeruleus are not known for their abundant dopaminergic innervation so we tested whether these could belong to other classes of neurons expressing TH. Co-immunolabeling of TH with Norepinephrine Transporter (NET) revealed that nearly all the spheroids outside the striatal regions were NET+ (Figure 5J). In agreement with its known localization, NET signal surrounded the TH+ signal indicating a membrane bound morphology at the spheroids (Figure 5J) (Mandela and Ordway, 2006). As mentioned earlier, we first observed these spheroids appear at 3-months of age and progress longitudinally (Figure S2). We did not notice differences in the number of these spheroids between hemizygous males and homozygous females, but heterozygous females appeared to have fewer spheroids than the full knockout animals (Figure 5 H-I)). Together, these results demonstrate a novel finding that both dopaminergic and noradrenergic neurons of a BPAN mouse model bear defects in their distal axons. This is consistent with motor defects and sleep disturbances that are commonly observed in BPAN patients.

### 3.8 Purkinje neurons and interneurons of Wdr45 c52C>T mice also develop spheroids

We further tested whether TH+ neurons are the only neurons affected in our BPAN mouse model. Previously, Zhao et al. reported very large Calbindin (CALB1)-positive spheroids in the deep cerebellar nuclei (DCN), starting at 1.5-month of age (Zhao et al., 2015). Most probably, these spheroids belonged to the axonal projections of cerebellar Purkinje neurons (Baumel et al., 2009). Therefore, we probed for the presence of these spheroids in our *Wdr45 c52C>T* model. As soon as P14 and in all of the *Wdr45 c52C>T* animals, CALB1 signal in the Deep Cerebellar Nuclei (DCN) revealed massive spheroids, which were much larger than in any other brain region (Figure 6B). These spheroids were positive for the presynaptic vesicle protein Synaptophysin (SYP), supporting the idea that these arise from axonal terminals and presynaptic compartments (Figure 6F). In addition to the DCN, we observed CALB1 positive spheroids throughout the brain at 3-months, including in the cortex and the hippocampus. This suggested that other subclasses of neurons may be affected. Since the majority of neurons expressing CALB1 are interneurons, we tested for the presence of spheroids using immunolabelling of Parvalbumin (PV) and Calretinin (CALB2). We observed similar spheroid structures in all mutant animals starting at 3-months and PV spheroids appeared as soon as 1-month, though the number of PV positive neurons is unchanged at 3-months (Figure 6C-D). Of note, the distribution of PV+ spheroids did not match that of the PV+ cell bodies, further hinting that PV+ spheroids arise from specific distal regions, not in the vicinity of cell bodies (Figure 6E). For all the markers we tested, spheroids were also present in heterozygous mice albeit with a lower density (Figure 6B). We further found similar results in CALB2 positive neurons (data not shown). Together, these data show that spheroids in WDR45 knockout animals are not exclusive to one type of neuron or within one particular area of the brain.

**Figure 6:**
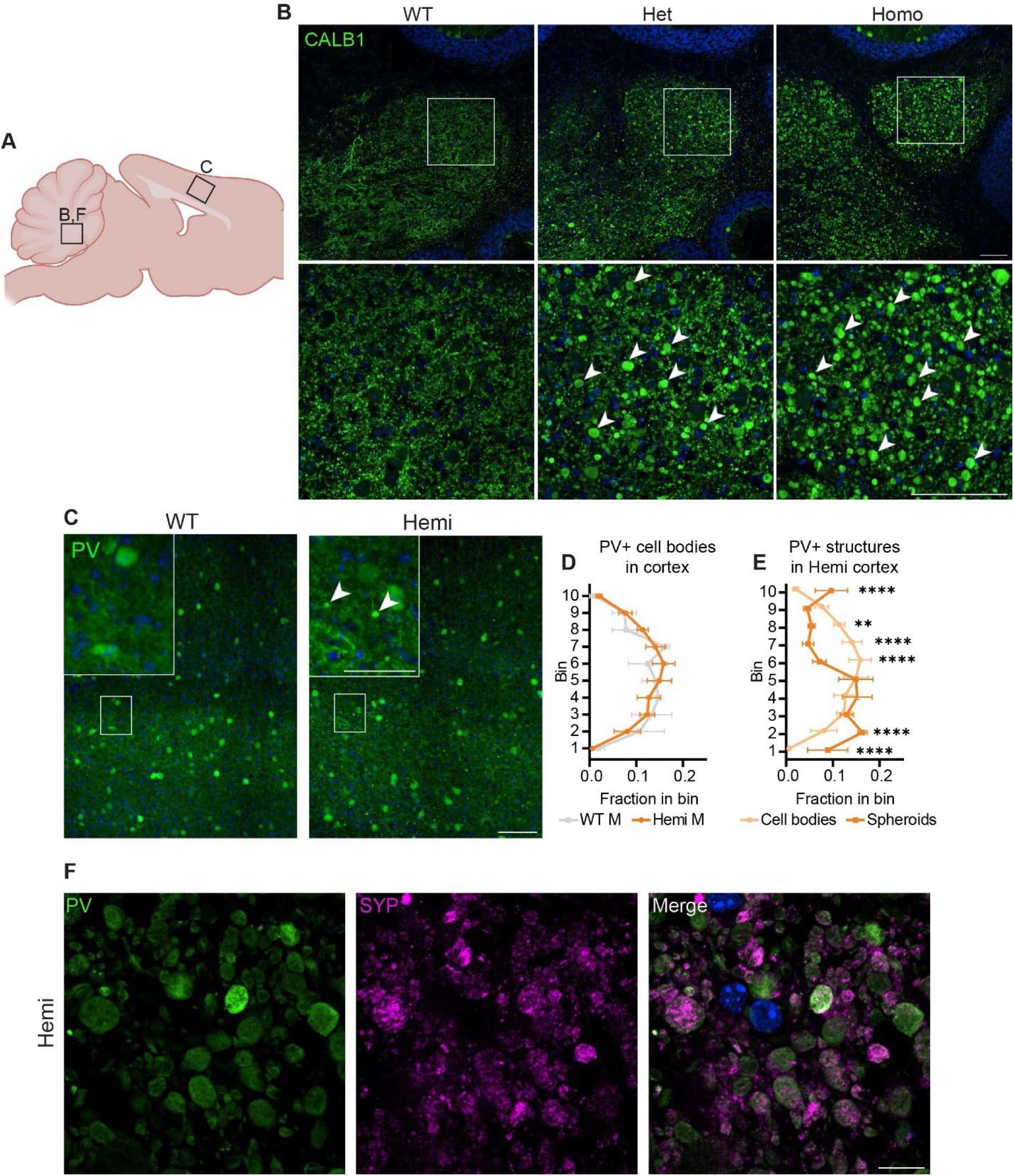
Spheroids are seen throughout the brain and in multiple neuronal sub-types in *Wdr45* c52C>T animals. Diagram of regions imaged (A). Calbindin (CALB1) positive swollen spheroids are apparent in the Deep Cerebellar Nuclei of *Wdr45* c52C>T animals at 1-month of age with (B). Parvalbumin (PV) labeling in the cortex of 3-month-old animals (C) showed no difference in number of Parvalbumin positive cell bodies or layer location of these cells between genotypes (D). Though, spheroids observed in the cortex of *Wdr45* c52C>T animals occupy different layer positions than the cells bodies (E). Colabeling in the DCN of PV (Green) and Synaptophysin (SYP) (Magenta) show accumulation of synaptic vesicles in these spheroids (F). Scale bars: B-C, F = 100µm. D, E used Two-way ANOVA with Holm-Šídák post hoc test, N=3-4 animals for each group. Mean ± SEM: * = p<0.05, ** = p<0.01, *** = p<0.001, **** = p<0.0001.

### 3.9 Wdr45 c52 C>T mutation leads to glial inflammation in mice

Astrocytic activation is a common marker of neuropathology and can indicate the progression of neurodegenerative diseases (Ben Haim et al., 2015, Jurga et al., 2020, Cain et al., 2019). We investigated GFAP levels, a marker for astrocytes that is overexpressed during glial activation, finding elevated immunostaining in the cortex of hemizygous males and homozygous females at 3-months of age (Figure 7C-E). We observed an increase in the heterozygous animals though deviation from wildtype levels was only significant at 12-months of age and is lower than the homozygous females at all timepoints (Figure 7D). We additionally saw an increase in GFAP signal localized to DCN in all mutant animals at 1-month through until 12-months (Figure 7F-H). Next, we labeled for the microglia protein CD68, which is upregulated during microglial activation. Immunolabeling for this protein showed increased staining in the cortex of homozygous female mice at 6-months and 12-months of age (Figure 7J-K). When we analyzed the CD68 signal in the DCN, hemizygous males and homozygous females showed increased signal at 1-month of age (Figure 7M-O). Together these data show an elevation in glial activation is consistent with age and disease progression in these mice with increase CD68 signal and GFAP positive astrocytes showing a robust increase in these brain regions.

**Figure 7:**
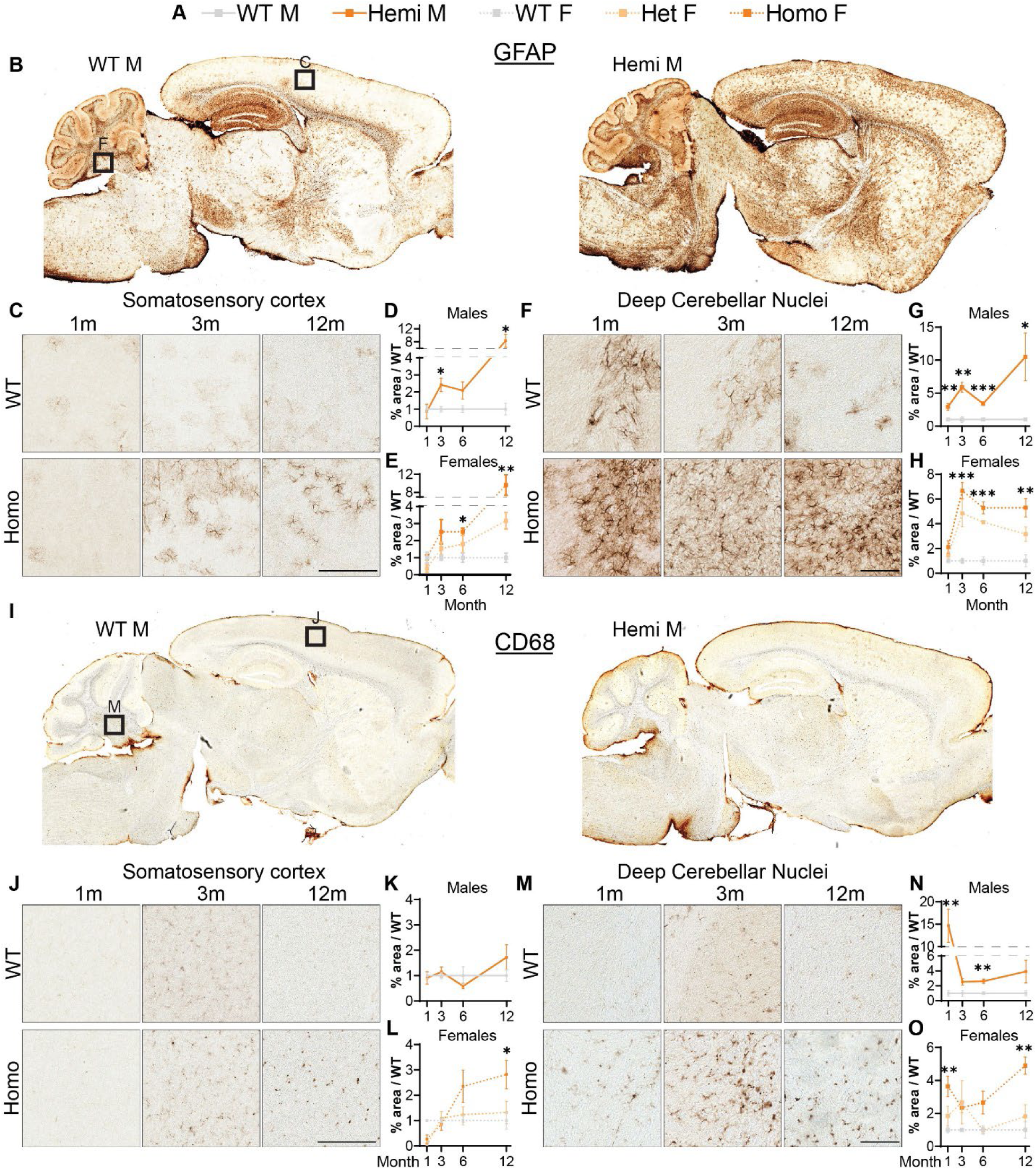
Glial activation markers are increased in *Wdr45* c52C>T animals. Genotype key for line graphs (A). Full sagittal scans of GFAP (B) and CD68 (I) at 12-months of age. Quantifications are area of immunopositive signal in each imaged field normalized to average values from sex and age matched wildtype animals. GFAP immunostaining was increased in the cortex (C,D,E) and DCN (F,G,H) of *Wdr45* c52C>T animals. CD68 immunostaining in the cortex (J,K,L) and DCN (M,N,O) also had increased immunopositivity in *Wdr45* c52C>T animals though these increases were much less consistent across timepoints. Scale bars: C,F,J,M = 100µm. For each timepoint D,G,K,N use T-Test and E,H,L,O use One-way ANOVA with Holm-Šídák post hoc test. N=3-4 animals for each group. Mean ± SEM: * = p<0.05, ** = p<0.01, *** = p<0.001, **** = p<0.0001.

### 3.10 Wdr45 c52 C>T mutant mice display altered autophagy, synaptic and iron regulating pathways

We next wanted to uncover pathways disrupted by loss of WDR45 that could be contributing to the phenotypes we described above. As an unbiased approach to identify novel pathways contributing to BPAN, we performed next-generation RNA sequencing on cortical samples in hemizygous and wildtype 3-month-old male littermates. This analysis revealed many differentially expressed genes between genotypes. Interestingly, we saw over three times as many genes downregulated (412 genes) compared to upregulated (128 genes) in the hemizygous mutant animals (Figure 8B). Gene ontology and pathway analysis was performed on the up and downregulated genes separately (Figure 8C-E). Notably, many synaptic, neuron projection, and neurotransmitter transport genes were downregulated in hemizygous cortices which may reflect the neuropathology we observed at this timepoint, and specifically axonal distress. We also saw increased expression of genes associated with ferroptosis, mitochondrial regulation, and membrane bound organelles which align with previous work related to WDR45 in cell culture models (Bakula et al., 2017, Ingrassia et al., 2017, Seibler et al., 2018, Cong et al., 2021, Zhu et al., 2024). Interestingly, the most significant differences in gene expression are seen with increased expression of the iron regulating gene Transferrin Receptor 1 (*Tfrc*) and down regulation of the transcript encoding the ER-associated, calcium-binding protein Calreticulin (*Calr*) (Figure 7B). This also agrees with cell models of WDR45 loss which show accumulation of TFRC, though this is the first time this has been shown at the transcriptional level and in an animal model of BPAN. To confirm the significance of the altered transcripts, we then probed for levels of the encoded proteins in cortical samples via western blotting. We found that both TFRC and CALR were not altered at 1-month in Wdr45 c52C>T mice. However, both proteins appeared elevated later in the life of these animals (Figure 9A-J). Further, when we probed for CALR using immunofluorescence, we found large areas of accumulation that appeared to coincide in size and position with the some of the swollen axonal structures we observed throughout the brain (Figure 9K). Co-immunolabeling of CALR together with glial markers GFAP (astrocytes) or IBA1 (microglia) showed no overlap between these markers and the CALR+ accumulations (Figure S3C-D).

**Figure 8:**
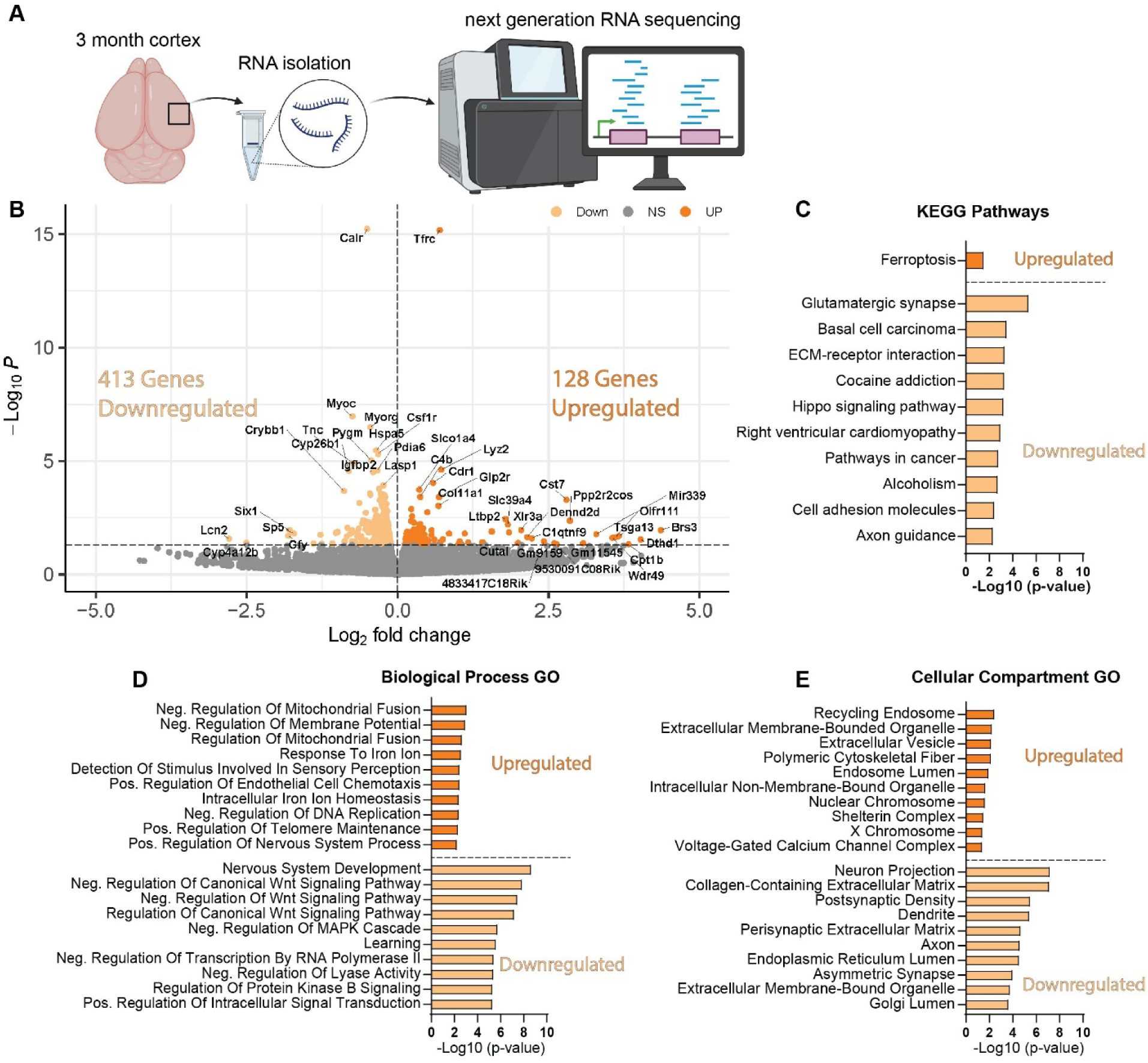
*Wdr45* c52C>T animals have altered synaptic, neurite associated, and ferroptosis related genes. RNA sequencing was performed on cortical samples from 3-month-old male animals (A). There were three times more downregulated DEGs than upregulated in Wdr45 c52C>T animals (B). Pathway and Gene ontology analysis was performed on up- and down-regulated DEGs separately with the top 10 terms (defined by highest -Log(p-value ≤ 0.05)) shown here (C-E). Pathway analysis of the DEGs indicate upregulation of ferroptosis related genes and down-regulation of synaptic genes (C). Gene Ontology analysis display altered expression of metabolism, iron regulation, synapse, and axon genes (D,E)

**Figure 9:**
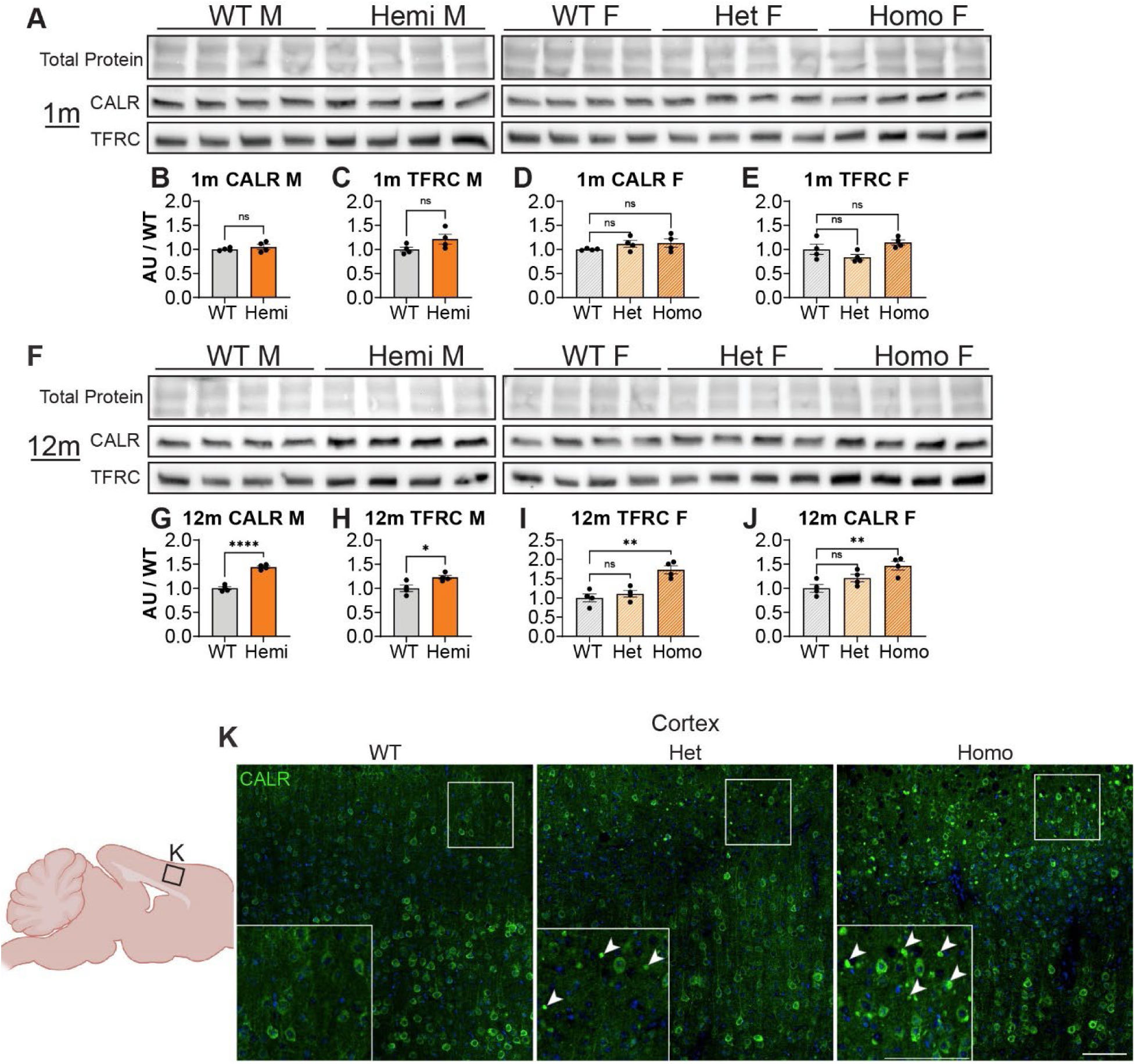
*Wdr45* c52C>T animals display accumulations of CALR in the cortex. Western blotting was performed on cortical samples from 1-month (A) and 12-month (F) animals. In *Wdr45* c52C>T animals CALR (B,D,G,I) and TFRC (C,E,H,J) were unchanged at 1 month and elevated at 12 months in Homozygous and Hemizygous animals. CALR and TFRC signal was normalized to total protein for each lane. (note these were run on the same gel and therefore have the same total protein as figure 1) Labeling of the Endoplasmic Reticulum resident protein Calreticulin (CALR) in the cortex reveals immunopositive puncta similar to the spheroids observed in the cortex in *Wdr45* c52C>T animals (K). All analyses were normalized to the average wildtype value for each experiment. Scale bars: K and K inset = 100µm. Analysis of males used T-Testing and analysis of females used One-way ANOVA with Holm-Šídák post hoc test. N=4 animals for each group. Mean ± SEM: * = p<0.05, ** = p<0.01, *** = p<0.001, **** = p<0.0001.

## 4 Discussion

In this article, we report a novel BPAN mouse model mimicking a patient variant in *Wdr45* and exhibiting neurologic symptoms in line with those observed in BPAN patients. This includes motor decline and intellectual deficits that arise at timepoints similar to BPAN progression. We observe glial inflammation and swollen structures in the distal axons of TH+, NET+, CALB1+, and PV+ neurons starting between 1 and 3-months of age in numerous brain regions. In the DCN, very large structures were observed as soon as P14. Thus, our novel model displays many phenotypes earlier, and in a wider variety of neuron subtypes and brain regions than previously reported. Importantly, we see many of these phenotypes occurring in heterozygous female animals, genetically reflecting a majority of the BPAN patient population. These animals often present with intermediate phenotypes when compared to homozygous and wildtype animals, though neurological pathology and dysfunction is clear. Not only will this novel model be helpful for our understanding of the disease, but it also provides numerous outcome measures that will be invaluable for therapeutic development. Finally, we also demonstrated that this model can be used to discover novel molecular pathways associated with this disease which reveal underlying neuronal biology.

### 4.1 Behavioral deficits observed in the C52c>t model

Using a suite of behavioral assays, we report neurologic deficits in *Wdr45* C>T mice that align with BPAN symptoms. We report these mice display hyperactivity in the forceplate assay and reduced vocalizations which have been previously associated with attention deficit and autism-like behaviors in other mouse models (Moy et al., 2009, Shekel et al., 2021). This reflects the behavioral issues seen in early-stage BPAN (Gavazzi et al., 2022). Our model also displays reduced performance in the rotarod and poleclimb assays, as well as reduced efficiency in the Morris water maze assay. These reflect motor decline and intellectual deficits, which coincide with the degenerative stage of the disease (Hayflick et al., 2013).

The phenotypic characterization of the model should be further refined in future studies. Most of the assays used here started at 3-months of age, with phenotypes already apparent at this timepoint. It will be important to clarify whether these phenotypes develop earlier. Given how early we see a vocalization deficit in our mice, it is likely that neurological deficits occur prior to the timepoints tested in the other assays. Moreover, some phenotypes in BPAN patients are lacking in our mouse models including spontaneous seizures and premature death (Wan et al., 2020, Zhao et al., 2015, Biagosch et al., 2021, Wang et al., 2024). These differences between mouse models and humans are not uncommon in neurodegenerative diseases, and may reflect fundamental differences in murine and human biology (Sasaguri et al., 2017). With that said, our current model provides robust readouts to explore how the underlying cellular and molecular mechanisms altered by *WDR45* loss leads to disease.

### 4.2 Impact on neurons and synaptic structures

We outline a clear phenotype whereby progressive, dramatic motor dysfunction coincides with terminal axon swelling in Purkinje neurons and dopaminergic neurons of the nigral-striatal pathway. A recent publication showed that selective ablation of WDR45 in DAT-expressing neurons leads to a similar swollen axon phenotype in the striatum, albeit at a much later timepoint (Wang et al., 2024). Whether this delayed phenotype is due to the selective nature of the ablation or that the recombination was not induced until the mice were 8 weeks of age, is unclear. Though, it does provide evidence that loss of *Wdr45* in neurons alone is sufficient to cause the axonal spheroids we observe. This suggests that WDR45 plays an essential role in neuron biology to maintain distal axon integrity likely contributing to motor and cognitive function. This is further supported by our RNA sequencing results which indicate reduced transcripts encoding synaptic and axonal proteins in the cortex of WDR45 deficient mice. Our data indicates this neuronal distress is widespread showing spheroids in multiple neuronal sub-types and in several brain regions. Importantly, work is still needed to uncover whether these spheroids appear in all neuronal subtypes, how these swellings form in WDR45 deficient mice, and how this directly impacts broader neuronal function.

Of note, we did not observe any iron accumulation, a common finding in BPAN patients, which is consistent with other mouse models lacking *Wdr45* expression at the timepoints observed (data not shown) (Wan et al., 2020, Zhao et al., 2015). This may be due to the lack of sensitivity of the Prussian Blue assay or this may relate to species-specific differences. None-the-less, we report several overlapping phenotypes that suggest this model will be useful for understanding the disease. Indeed, the swellings we report appear similar to the axonal spheroids that have been reported in BPAN patients and other neurodegenerative disorders (Paudel et al., 2015, Yong et al., 2021).

### 4.3 Insights on the molecular pathways involved in BPAN

It is currently unclear what role WDR45 plays that, when lost, leads to the neurological distress we see in patients and mice. WDR45 is one of a family of WIPI proteins which are mammalian orthologs of the yeast protein atg18. Atg18 plays an important part in the formation of the autophagosome and progression of bulk autophagy in yeast cells (Proikas-Cezanne et al., 2015). Some of the first research into WDR45 function in mammalian cells established an interaction with ATG2 and binding of WDR45 to PI3P rich regions of membrane, and particularly the developing autophagosome (Proikas-Cezanne et al., 2015). Though, neuroblastoma cells lacking WDR45 expression do not have a reduced number of autophagosomes and a recent publication showed autophagy induction does not rescue retinal degeneration in WDR45 deficient zebra fish (Ji et al., 2020, Zhu et al., 2024). This suggests that WDR45 associated neurodegeneration may be more complicated than solely disrupted autophagy regulation.

What other roles of WDR45 could lead to the phenotypes we observed? Mitochondrial dysfunction, disruptions in iron metabolism, lipid peroxidation, ER stress, and lipid processing issues have all been observed in WDR45 deficient cell and animal models (Cong et al., 2021, Biagosch et al., 2021, Wan et al., 2020, Zhao et al., 2015, Hinarejos et al., 2020, Ingrassia et al., 2017). Our data suggests that many of these cellular mechanisms may be disrupted in the brain of *Wdr45* mutant animals, including mitochondrial regulation, iron response, and vesicular regulation. It is unclear what role WDR45 plays in these processes or which of the pathways it touches are critical for the neurological dysfunction we see in BPAN patients. Importantly, many of these functions have not been explored in the nervous system. Providing robust behavioral, cellular, and molecular phenotypes, we predict our model will be critical for exploring WDR45 functions in a more relevant cellular and histological context, and how alteration of these functions leads to BPAN.

## Data Availability Statement

The original contributions presented in the study are included in the article/Supplementary Material, further inquiries can be directed to the corresponding author.

## Ethics Statement

The animal study was reviewed and approved by Sanford Research IACUC.

## Author Contributions

The project was conceived by BLM, JMW, and LJP. BLM and LJP designed experiments. Experiments were performed by BLM, KSK, MJR, PRP, ACE, JMH, ERA, and LJP. The manuscript was prepared by BLM and LJP. BLM, KSK, MJR, PRP, ACE, JMH, ERA, JMW, and LJP reviewed and edited the manuscript. All authors contributed to the article and approved the submitted version.

## Acknowledgments

The authors thank Dr. Jon Brudvig for his input and guidance in the development of this project. Cartoons created with BioRender.com.

---

**Figure S1:**
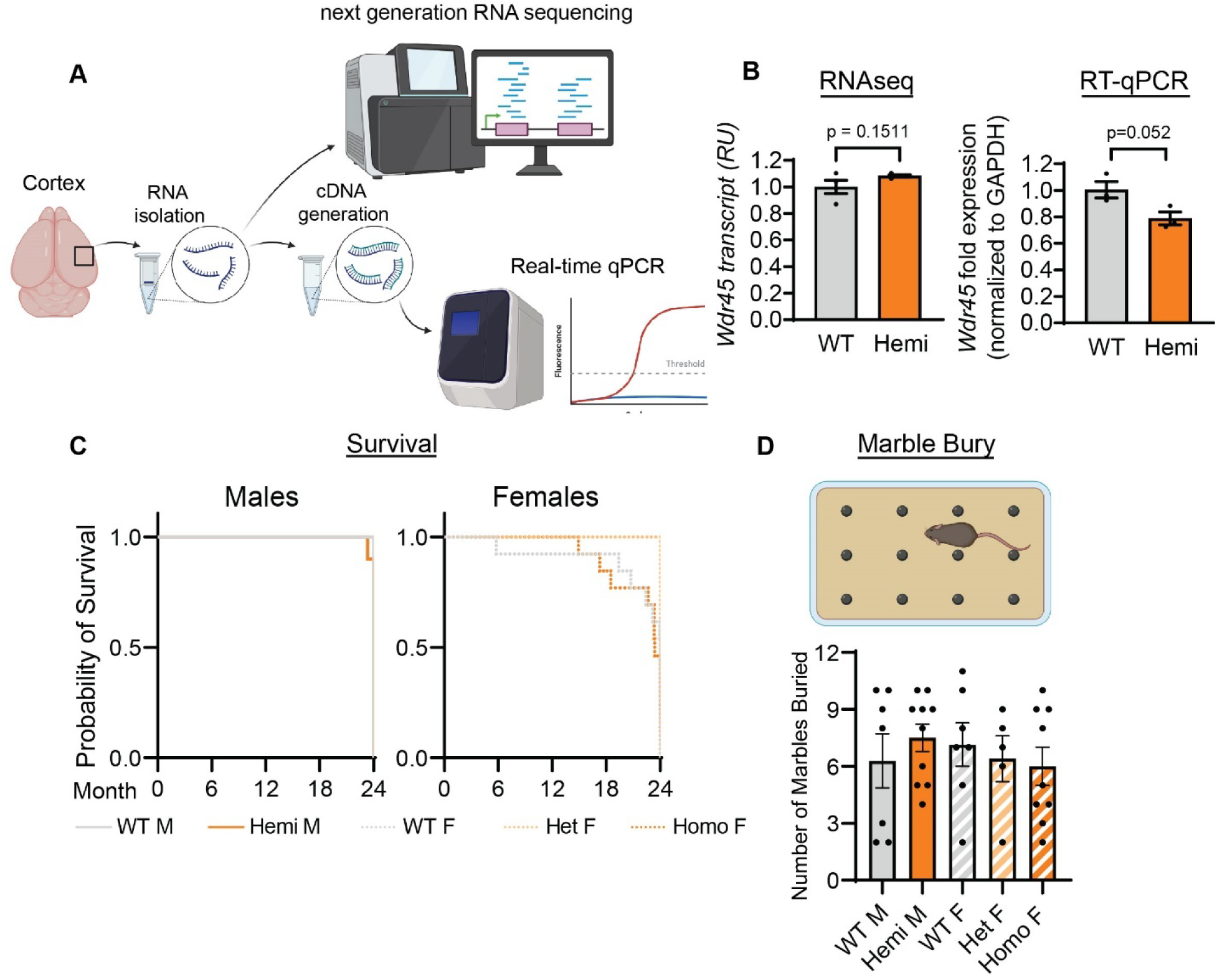
*Wdr45* c52C>T animals have unchanged Wdr45 transcript but exhibit unaltered survival and anxiety phenotypes. *Wdr45* c52C>T animals have unchanged transcript abundance in the cortex at 3 months of age (B). *Wdr45* c52C>T animals showed unchanged survival through two years of age (B). Additionally, no difference was seen between genotypes in the marble bury test performed at 3 months of age (D). For D: Testing was performed using Mantel Cox test on males and females separately. For B: Males were tested using a T-test. Females were tested using a one-way ANOVA with Holm-Šídák post hoc test. N= 8-14 per group. Mean ± SEM

**Figure S2:**
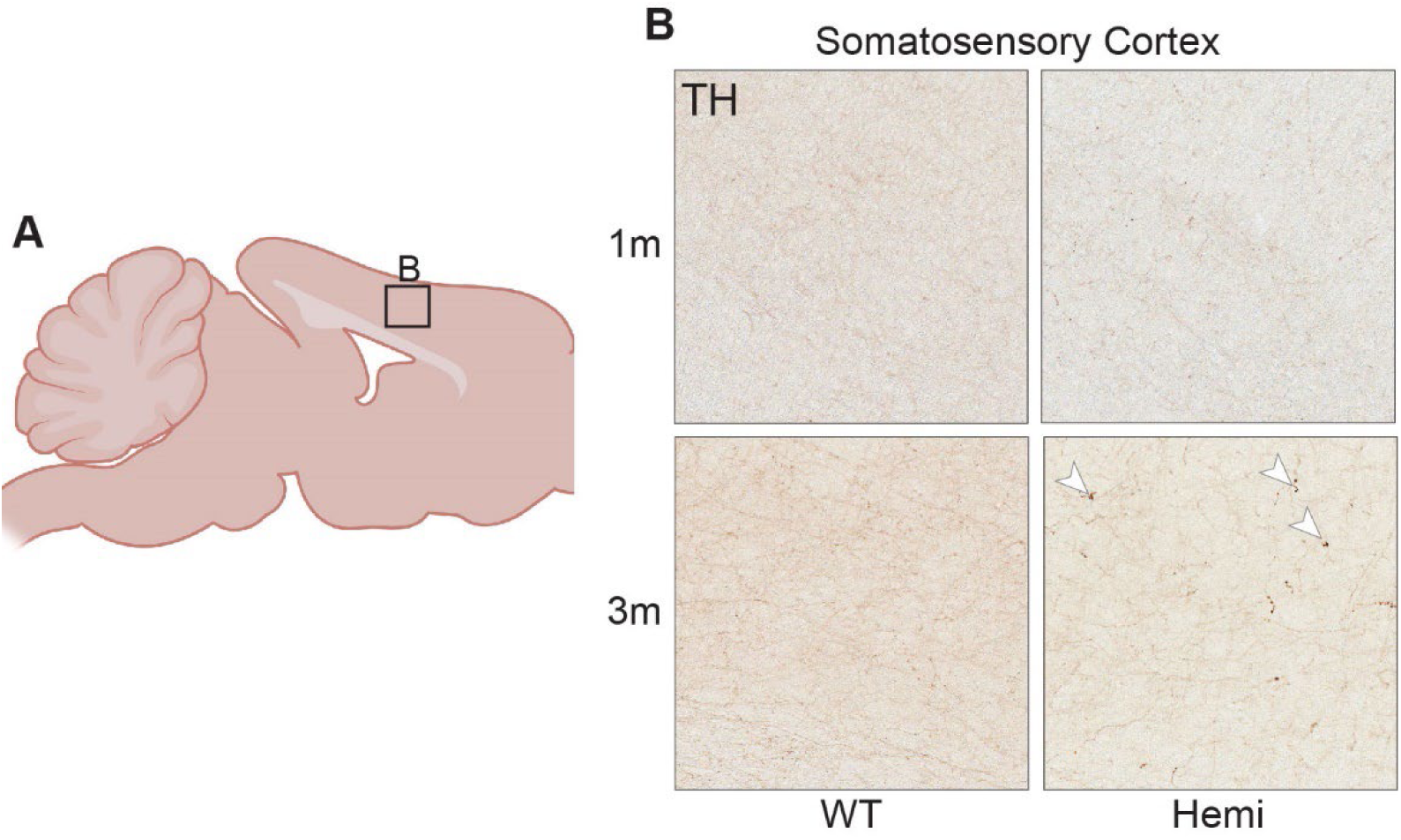
TH positive spheroids appear at 3 months of age in WDR45 c52C>T mice. Diagram for area imaged (A). TH immunolabeling at earlier timepoints indicate absence of these structures at 1 month of age, and presence at 3 months of age in *Wdr45 c52C>T* animals (B).

**Figure S3:**
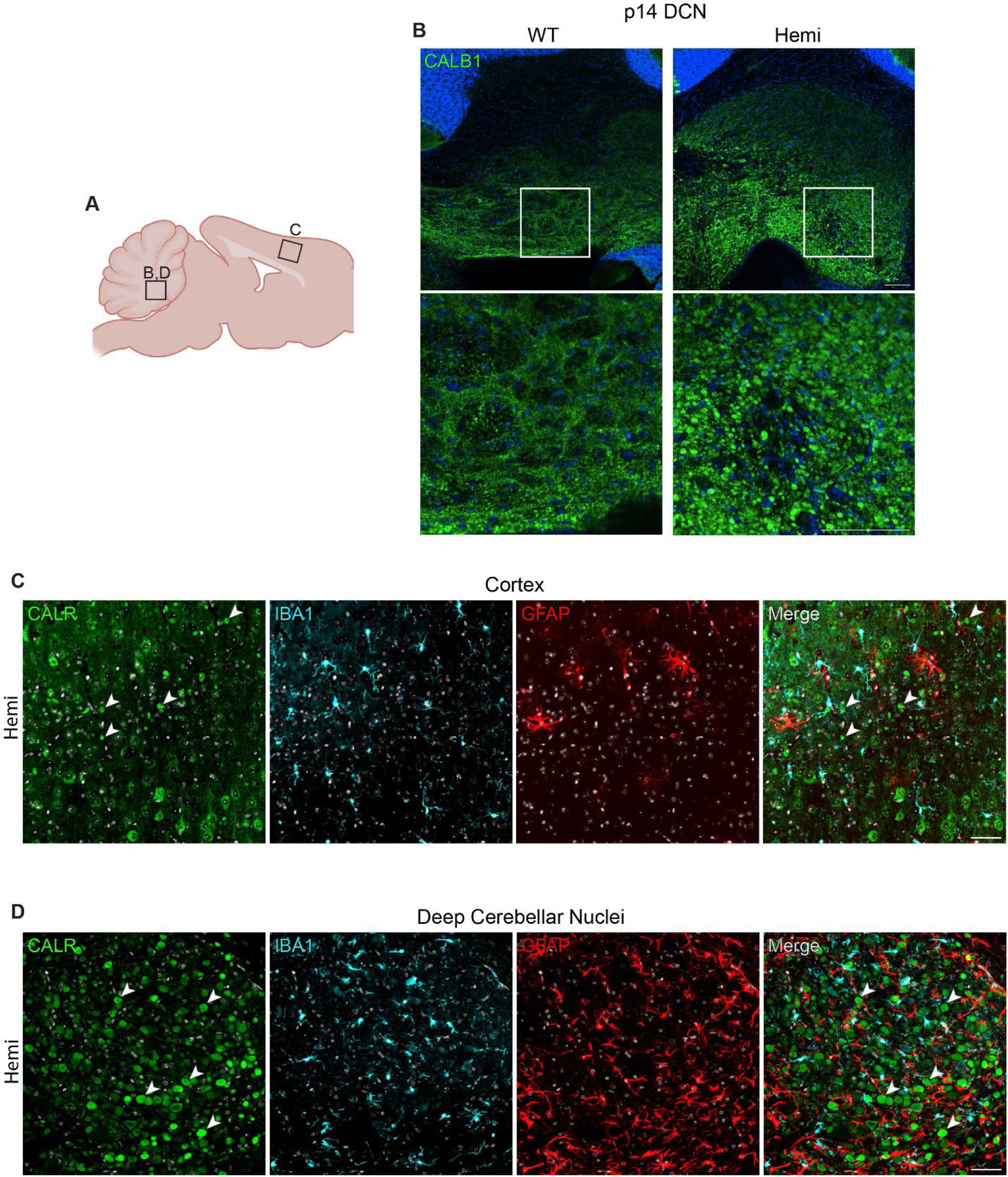
DCN spheroids appear at P14 and CALR puncta are distinct from glial cells in *Wdr45* c52C>T mice. Diagram for areas imaged (A). CALBINDIN (Calb1 - Green) labeling in the DCN at P14 shows spheroid accumulation at this early timepoint (B). CALR (Green) puncta do not colocalize with microglia (IBA1 - Cyan) or astrocyte (GFAP - Red) markers (C,D) at 6 months of age. Scale bars: B-D = 100µm

